# Accessible and reproducible mass spectrometry imaging data analysis in Galaxy

**DOI:** 10.1101/628719

**Authors:** Melanie Christine Föll, Lennart Moritz, Thomas Wollmann, Maren Nicole Stillger, Niklas Vockert, Martin Werner, Peter Bronsert, Karl Rohr, Björn Andreas Grüning, Oliver Schilling

## Abstract

**Background:** Mass spectrometry imaging is increasingly used in biological and translational research as it has the ability to determine the spatial distribution of hundreds of analytes in a sample. Being at the interface of proteomics/metabolomics and imaging, the acquired data sets are large and complex and often analyzed with proprietary software or in-house scripts, which hinder reproducibility. Open source software solutions that enable reproducible data analysis often require programming skills and are therefore not accessible to many MSI researchers.

**Findings:** We have integrated 18 dedicated mass spectrometry imaging tools into the Galaxy framework to allow accessible, reproducible, and transparent data analysis. Our tools are based on Cardinal, MALDIquant, and scikit-image and enable all major MSI analysis steps such as quality control, visualization, preprocessing, statistical analysis, and image co-registration. Further, we created hands-on training material for use cases in proteomics and metabolomics. To demonstrate the utility of our tools, we re-analyzed a publicly available N-linked glycan imaging dataset. By providing the entire analysis history online, we highlight how the Galaxy framework fosters transparent and reproducible research.

**Conclusion:** The Galaxy framework has emerged as a powerful analysis platform for the analysis of MSI data with ease of use and access together with high levels of reproducibility and transparency.

## Findings

### Background

Mass spectrometry imaging (MSI) is increasingly used for a broad range of biological and clinical applications as it allows the simultaneous measurement of hundreds of analytes and their spatial distribution. The versatility of MSI is based on its ability to measure many different kinds of molecules such as peptides, metabolites or chemical compounds in a large variety of samples such as cells, tissues, fingerprints or human made materials [1–5]. Depending on the sample, the analyte of interest and the application, different mass spectrometers are used [6]. The most common ionization sources are MALDI (Matrix Assisted Laser Desorption/Ionization), DESI (Desorption Electrospray Ionization) and SIMS (Secondary Ion Mass Spectrometry). Typical mass analyzers are time of flight (TOF) devices and ion traps.

Due to the variety of samples, analytes, and mass spectrometers, MSI is suitable for highly diverse use cases ranging from plant research, to (pre-)clinical, pharmacologic studies, and forensic investigations [2, 7–9]. On the other hand, the variety of research fields hinders harmonization and standardization of MSI protocols. Recently efforts were started to develop optimized sample preparation protocols and show their reproducibility in multicenter studies [10–13]. In contrast, efforts to make data analysis standardized and reproducible are in its infancy.

Reproducibility of MSI data analyses is hindered by the common use of software with restricted access such as proprietary software, license requiring software, or unpublished in-house scripts [14]. Open source software has the potential to advance accessibility and reproducibility issues in data analysis but requires complete reporting of software versions and parameters, which is not yet routine in MSI [15–17].

At the same time, the introduction of the open standard file format imzML has opened new avenues to the community and an increasing number of open source software tools are emerging [18]. Yet, many of these tools necessitate steep learning curves, in some cases even requiring programming knowledge to make use of their full range of functions [19–23].

To overcome problems with accessibility of software and computing resources, standardization, and reproducibility, we developed MSI data analysis tools for the Galaxy framework that are based on the open source software suites Cardinal, MALDIquant, and scikit-image [20, 21, 24]. Galaxy is an open source computational platform for biomedical research that was developed to support researchers without programming skills with the analysis of large data sets, e.g. in the field of next generation sequencing. Galaxy is used by hundred thousands of researchers and provides thousands of different tools for many different scientific fields [25].

### Aims

With the present publication, we aim to raise awareness within the MSI community for the advantages being offered by the Galaxy framework with regard to standardized and reproducible data analysis pipelines. Secondly, we present newly developed Galaxy tools and offer them to the MSI community through the graphical front-end and “drag-and-drop” workflows of the Galaxy framework. Thirdly, we apply the MSI Galaxy tools to a publicly available dataset to study N-glycan identity and distribution in murine kidney specimens in order to demonstrate usage of a Galaxy-based MSI analysis pipeline that facilitates standardization and reproducibility and is compatible with the principles of FAIR (findable, accessible, interoperable, and re-usable) data and MIAPE (minimum information about a proteomics experiment) [26, 27].

#### The Galaxy framework for flexible and reproducible data analysis

In essence, the Galaxy framework is characterized by four hallmarks: (1) usage of a graphical front-end that is web browser based, hence alleviating the need for advanced IT skills or the requirement to locally install and maintain software tools; (2) access to large-scale computational resources for academic users; (3) provenance tracking and full version control, including the ability to switch between software and tool version and to publish complete analysis, thus enabling full reproducibility; (4) access to a vast array of open-source tools with the ability of seamless passing data from one tool to another, thus generating added value by interoperability.

Multiple Galaxy servers on essentially every continent provide access to large computing resources, data storing capabilities, and hundreds of pre-installed tools for a broad range of data analysis applications through a web browser based graphical user interface [28–30]. Additionally, there are more than hundred public Galaxy servers available that offer more specific tools for niche application areas. For local usage, Galaxy can be installed on any computer ranging from private laptops to high computing clusters. So-called “containers” facilitate a fully functional one-click installation independent of the operating system. Hence, local Galaxy serves are easily deployed even in “private” network situations in which these servers remain invisible and inaccessible to outside users. This ability empowers Galaxy for the analysis of sensitive and protected data, e.g. in a clinical setting.

In the Galaxy framework, data analysis information is stored alongside the results of each analysis step to ensure reproducibility and traceability of results. The information includes tool and software names and versions together with all parameters [31].

We propose that MSI research can greatly benefit from the possibility to privately or publicly share data analysis histories, workflows, and visualizations with collaboration partners or the entire scientific community, e.g. as online supplementary data for peer-reviewed publications. The latter step easily fulfills the criteria of the suggested MSI minimum reporting guidelines [6, 16].

The Galaxy framework is predestinated for the analysis of multi-omics studies as it facilitates the integration of software of different origin into one analysis [32, 33]. The possibility to seamlessly link tools of different origins has outstanding potential for MSI studies that often rely on different software platforms to analyze MSI data, additional MS/MS data (from liquid chromatography coupled tandem mass spectrometry), and (multimodal) imaging data. More than hundred tools for proteomic and metabolomics data analysis are readily available in Galaxy due to community driven efforts [34–38]. Increasing integration of MSI with other omics approaches such as genomics and transcriptomics is anticipated and the Galaxy framework offers a powerful and future-proof platform to tackle complex, interconnected data-driven experiments.

#### The newly available MSI toolset in the Galaxy framework

We have developed 18 Galaxy tools that are based on the commonly used open-source softwares Cardinal, MALDIquant, and scikit-image and enable all steps that commonly occur in MSI data analysis (Figure 1) [20, 21, 24]. In order to deeply integrate those tools into the Galaxy framework, we developed bioconda packages and biocontainers as well as a so-called ‘wrapper’ for each tool [31, 39]. The MSI tools consist of R scripts that were developed based on Cardinal and MALDIquant functionalities, extended for more analysis options and a consistent framework for input and output of metadata (Additional File 1). The image co-registration method uses scikit-image for image processing. All tools are deliberately build in a modular way to enable highly flexible analysis and to allow a multitude of additional functionalities by cleverly combining the MSI specific tools with already availably Galaxy tools.

**Figure 1:**
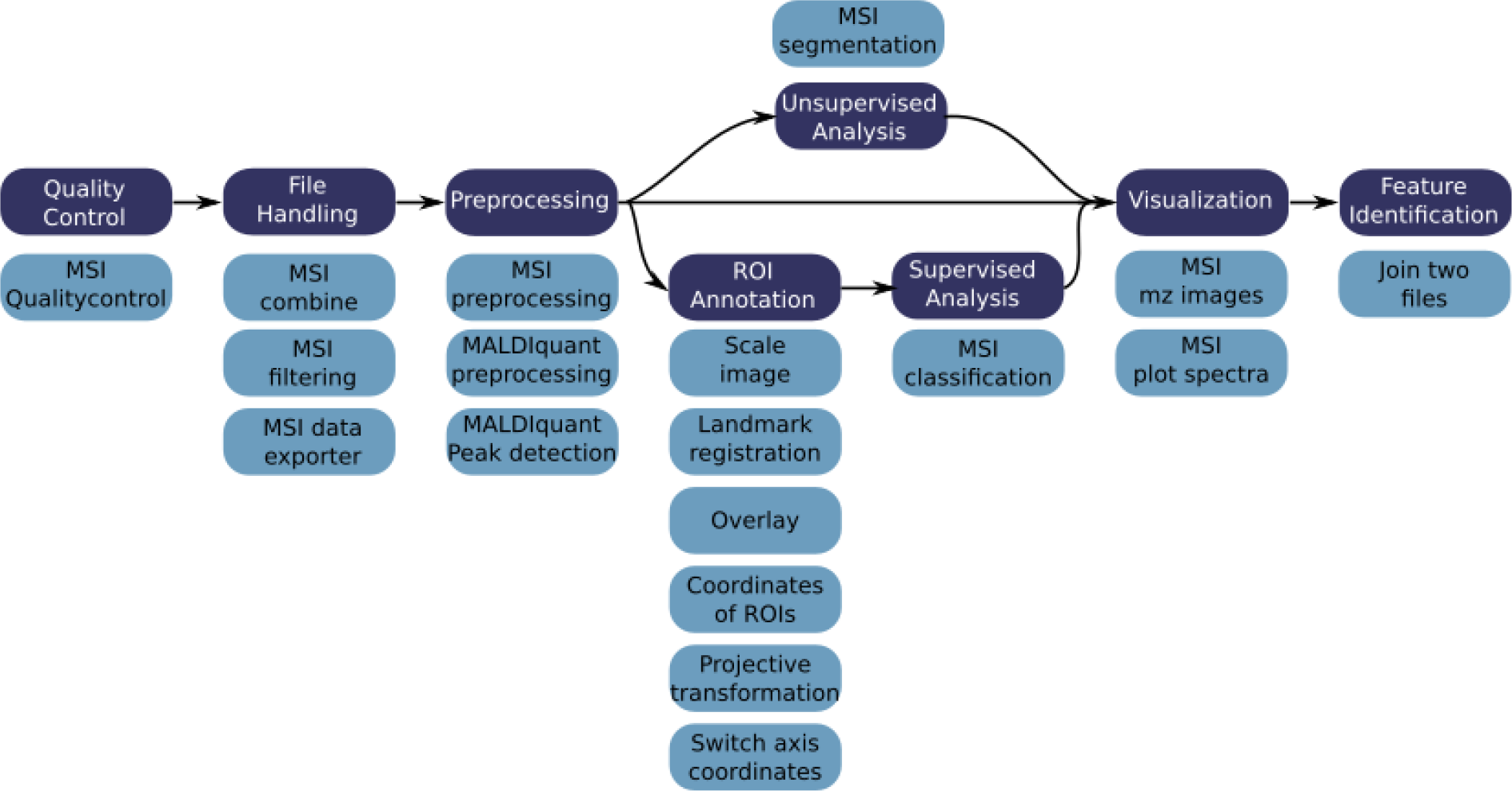
Typical MSI data analysis steps and associated Galaxy tools. Typical MSI data analysis step include Quality control, file handling, preprocessing, ROI annotation, supervised and unsupervised statistical analysis, visualizations and identification of features. Due to the variety of MSI applications, tools of all or only a few of these categories are used and the order of usage is highly flexible. To serve a broad range of data analysis tasks, we provide 18 tools that cover all common data analysis procedures and can be arbitrarily connected to allow customized analysis.

##### Data formats and data handling

We extended the Galaxy framework to support open and standardized MSI data files such as imzML, which is the default input format for the Galaxy MSI tools. Nowadays, the major mass spectrometer vendors directly support the imzML standard and several tools exist to convert different file formats to imzML [40]. Data can be easily uploaded to Galaxy via a web browser or via a built-in file transfer protocol (FTP) functionality. Intermediate result files can be further processed in the interactive environment that supports R Studio and Jupyter or downloaded for additional analysis outside of Galaxy [41].

To facilitate the parallel analysis of multiple files, the Galaxy framework offers so-called “file collections”. Numerous files can be represented in a file collection allowing simultaneous analysis of all files while the effort for the user is the same as for a single file. MSI meta data such as spectra annotations, calibrant m/z, and statistical results are stored as tab-separated values files, thus enabling processing by a plethora of tools both inside and outside the Galaxy framework. All graphical results of the MSI tools are stored as concise vector graphic PDF reports with publication-quality images.

##### Quality control and visualization

###### MSI Quality control

Quality control is an essential step in data analysis and should not only be used to judge the quality of the raw data but also to control processing steps such as smoothing, peak picking, and intensity normalization. Therefore, we have developed the ‘MSI Qualitycontrol’ tool that automatically generates a comprehensive pdf report with more than 30 different plots that enable a global view of all aspects of the MSI data including intensity distribution, m/z accuracy and segmentation maps. For example, spectra with bad quality, such as low total ion current or low number of peaks can be directly spotted in the quality report and subsequently be removed by applying the ‘MSI data exporter’ and ‘MSI filtering’ tools.

###### MSI mz image

The ‘MSI mz image’ tool allows to automatically generate a publication-quality pdf file with distribution heat maps for all m/z features provided in a tab-separated values file. Contrast enhancement and smoothing options are available as well as the possibility to overlay several m/z features in one image.

###### MSI plot spectra

The ‘MSI plot spectra’ tool displays multiple single or average mass spectra in a pdf file. Overlay of multiple single or averaged mass spectra with different colors in one plot is also possible.

The Galaxy framework offers various visualization options for tab-separated values files, including heatmaps, barplots, scatterplots, and histograms. This enables a quick visualization of the properties of tab-separated values files obtained during MSI analysis.

##### MSI file handling

A large variety of tools that allows for filtering, sorting, and manipulating of tab-separated values files is already available in Galaxy and can be integrated into the MSI data analysis. Some dedicated tools for imzML file handling were newly integrated into the Galaxy framework.

###### MSI combine

The ‘MSI combine’ tool allows combining several imzML files into a merged dataset. This is especially important to enable direct visual but also statistical comparison of MSI data that derived from multiple files. With the ‘MSI combine tool’, individual MSI datasets are either placed next to each other in a coordinate system or can be shifted in x or y direction in a user defined way. The output of the tool contains a single file with the combined MSI data and an additional tab-separated values file with spectra annotations, i.e. each spectrum is annotated with its original file name (before combination) and, if applicable, with previously defined annotations such as diagnosis, disease type, and other clinical parameters.

###### MSI filtering

The ‘MSI filtering’ tool provides options to filter m/z features and pixel (spectra) of interest, either by applying manual ranges (minimum and maximum m/z, spatial area as defined by x / y coordinates) or by keeping only m/z features or coordinates of pixels that are provided in a tab-separated values file. Unwanted m/z features such as pre-defined contaminant features can be removed within a preselected m/z tolerance.

###### MSI data exporter

The ‘MSI data exporter’ can export the spectra, intensity and m/z data of an imzML file together with their summarized properties into tab-separated values files.

##### Region of interest annotation

For supervised analysis, spatial regions of interest (ROI) can be defined. However, annotation of these ROIs is infeasible on the MSI images. Therefore, the ROIs are annotated on a photograph or histological image of the sample. We extended and developed six new Galaxy tools and combined them with existing tools into a workflow that enables co-registration of the real image (photograph or histological image), ROIs, and the MSI image by alignment using an affine transformation [42]. The transformation is estimated by a least-squares method using landmarks from both real and MSI image which are annotated outside Galaxy, for example, using the GNU Image Manipulation Program (GIMP) [43]. For more robust estimation of the transformation, Random sample consensus (RANSAC) is used on random subsets of landmark pairs.

We have also built automated workflows to convert annotation files from proprietary Bruker software (spotlist.txt and regions.xml) into annotation files that are compatible with the Galaxy MSI tools.

##### Preprocessing

Preprocessing of raw MSI spectra is performed to reduce data size and to remove noise, inaccuracies and biases to improve downstream analysis. Crucial steps are peak picking to reduce file size and remove noise features, intensity normalization to make spectra within and between different samples comparable, as well as m/z recalibration to improve comparability and identification of analytes.

###### MSI preprocessing

The ‘MSI preprocessing’ tool offers a multitude of algorithms that are useful to preprocess raw MSI data: intensity normalization to the total ion current (TIC), baseline removal, smoothing, peak picking, peak alignment, peak filtering, intensity transformation, binning and resampling.

###### MALDIquant preprocessing and MALDIquant peak detection

Both MALDIquant tools offer a multitude of preprocessing algorithms that complement those of the cardinal based MSI preprocessing tool such as m/z re-calibration, peak picking on average mass spectra and picking of monoisotopes.

##### Statistical analysis

A multitude of statistical analysis options for tab-separated values files is already available in Galaxy, the most MSI relevant tools are from the Workflow4metabolomics project and consist of unsupervised and supervised statistical analysis tools [44]. For specific purposes of spatially resolved MSI data analysis, we have integrated Cardinal’s powerful spatially aware statistical analysis options into the Galaxy framework.

###### MSI segmentation

The ‘MSI segmentation’ tool enables spatially aware unsupervised statistical analysis with principal component analysis, spatially aware k-means clustering and spatial shrunken centroids [45, 46].

###### MSI classification

The ‘MSI classification’ tool offers three options for spatially aware supervised statistical analysis: partial least square (discriminant analysis), orthogonal partial least squares (discriminant analysis), and spatial shrunken centroids [47].

##### Analyte identification

m/z determination on its own often remains insufficient to identify analytes. Compound fragmentation and tandem mass spectrometry are typically employed for compound identification by mass spectrometry. In MSI, the required local confinement of the mass spectrometry analysis severely limits the compound amounts that are available for fragmentation. Hence, direct on-target fragmentation is rarely employed in MSI. A common practice for compound identification includes a combinatorial approach in which LC-MS/MS data is used to identify the analytes while MSI analyses their spatial distribution. This approach requires assigning putative analyte information to m/z values within a given accuracy range.

###### Join two files on a column allowing a small difference

This newly developed tool allows for the matching of numeric columns of two tab-separated values files on the smallest distance that can be absolute or in ppm. This tool can be used to identify the m/z features of a tab-separated values files by matching them to already identified m/z features of another tab-separated values file (e.g. from a database or from an analysis workflow).

Community efforts such as Galaxy-M, Galaxy-P, Phenomenal, and Workflow4Metabolomics have led to a multitude of metabolomics and proteomics analysis tools available in Galaxy [34–38]. These tools allow analyzing additional tandem mass spectrometry data that is often acquired to aid identification of MSI m/z features. Databases to which the results can be matched, such as uniprot and lipidmaps, are directly available in Galaxy [48, 49]. The highly interdisciplinary and modular data analysis options in Galaxy render it a very powerful platform for MSI data analyses that are part of a multi-omics study.

#### Accessibility & training

All described tools are easily accessible and usable via the European Galaxy server [29]. Furthermore, all tools are deposited in the Galaxy Toolshed from where they can be easily installed into any other Galaxy instance [50]. We have developed bioconda packages and biocontainers that allow for version control and automated installation of all tool dependencies – those packages are also useful outside Galaxy to enhance reproducibility [31, 39]. For researchers that do not want to use publicly available Galaxy servers, we provide a pre-built Docker image that is easy to install independent of the operating system.

For a swift introduction into the analysis of MSI data in Galaxy, we have developed training material for metabolomics and proteomic use cases and deposited it to the central repository of the Galaxy Training Network [51, 52]. The training materials consist of a comprehensive collection of small example datasets, step-by-step explanations and workflows that enable any interested researcher in following the training and understanding it through active participation.

The first training explains data upload in Galaxy and describes the quality control of a mouse kidney tissue section in which peptides were imaged with an old MALDI-TOF [53]. The dataset contains peptide calibrants that allow the control of the digestion efficiency and m/z accuracy. Export of MSI data into tab-separated values files and further filtering of those files is explained as well.

The second training explains the examination of the spatial distribution of volatile organic compounds in a chili section. The training roughly follows the corresponding publication and explains how average mass spectra are plotted and only the relevant m/z range is kept, as well as how to automatically generate many m/z distribution maps and overlay several m/z feature maps [19].

The third training determines and identifies N-linked glycans in mouse kidney tissue sections with MALDI-TOF and additional LC-MS/MS data analysis [54, 55]. The training covers combining datasets, preprocessing as well as unsupervised and supervised statistical analysis to find potential N-linked glycans that have different abundances in the PNGase F treated kidney section compared to the kidney section that was treated with buffer only. The training further covers identification of the potential N-linked glycans by matching their m/z values to a list of N-linked glycan m/z that were identified by LC-MS/MS. The full dataset is used as a case study in the following section.

#### Case study

To exemplify the utility of our MSI tools we re-analyzed the N-glycan dataset that was recently made available by Gustafsson et al. via the PRIDE repository with accession PXD009808 [55, 56]. The aim of the study was to demonstrate that their automated sample preparation method for MALDI imaging of N-linked glycans successfully works on formalin-fixed paraffin-embedded (FFPE) murine kidney tissue [54]. PNGase F was printed on two FFPE murine kidney sections to release N-linked glycans from proteins while in a third section one part of the kidney was covered with N-glycan calibrants and another part with buffer to serve as a control. We downloaded all four imzML files (two treated kidneys, control and calibrants) from PRIDE and uploaded them with the composite upload function into Galaxy. To obtain an overview of the files we used the ‘MSI Qualitycontrol’ tool. We resampled the m/z axis, combined all files and run again the ‘MSI Qualitycontrol’ tool to directly compare the four subfiles. Next, we performed TIC normalization, smoothing and baseline removal. Spectra were aligned to the stable peaks that are present in at least 80 % of all spectra [57]. Spectra, in which less than two stable peaks could be aligned, were removed. This affected mainly spectra from the control file. Peak picking, detection of monoisotopic peaks and binning was performed on the average spectra of each subfile. The obtained m/z features were extracted with Cardinal’s ‘peaks’ algorithm from the normalized, smoothed, baseline removed and aligned file. Next, principal component analysis with four components was performed (Figure 2). To find potential N-linked glycans, the two treated tissues were compared to the control tissue with the supervised spatial shrunken centroids algorithm. Spatial shrunken centroids is a multivariate classification method that was specifically developed to account for the spatial structure of the data (Figure 3a) [45]. The supervised analysis provided us with 28 m/z features that discriminated between the two PNGase F treated kidneys and the control kidney with a spatial shrunken centroids p-value <0.05 and higher abundance in the treated kidneys. Matching those features that potentially are N-glycans to the m/z list of the original publication (Gustafsson et al., Supplementary Table S2) revealed the identity of 16 N-glycans with an average m/z error of 49 ppm (Table 1). Fifteen of those N-glycans match to the findings of the original publication. While we missed the 1647.635 m/z N-glycan, we found another N-glycan with 1542.62 m/z. The intensity distribution for four N-glycans on the TIC normalized dataset is depicted in Figure 3b-e and three of them are overlayed in Figure 3f.

**Figure 2:**
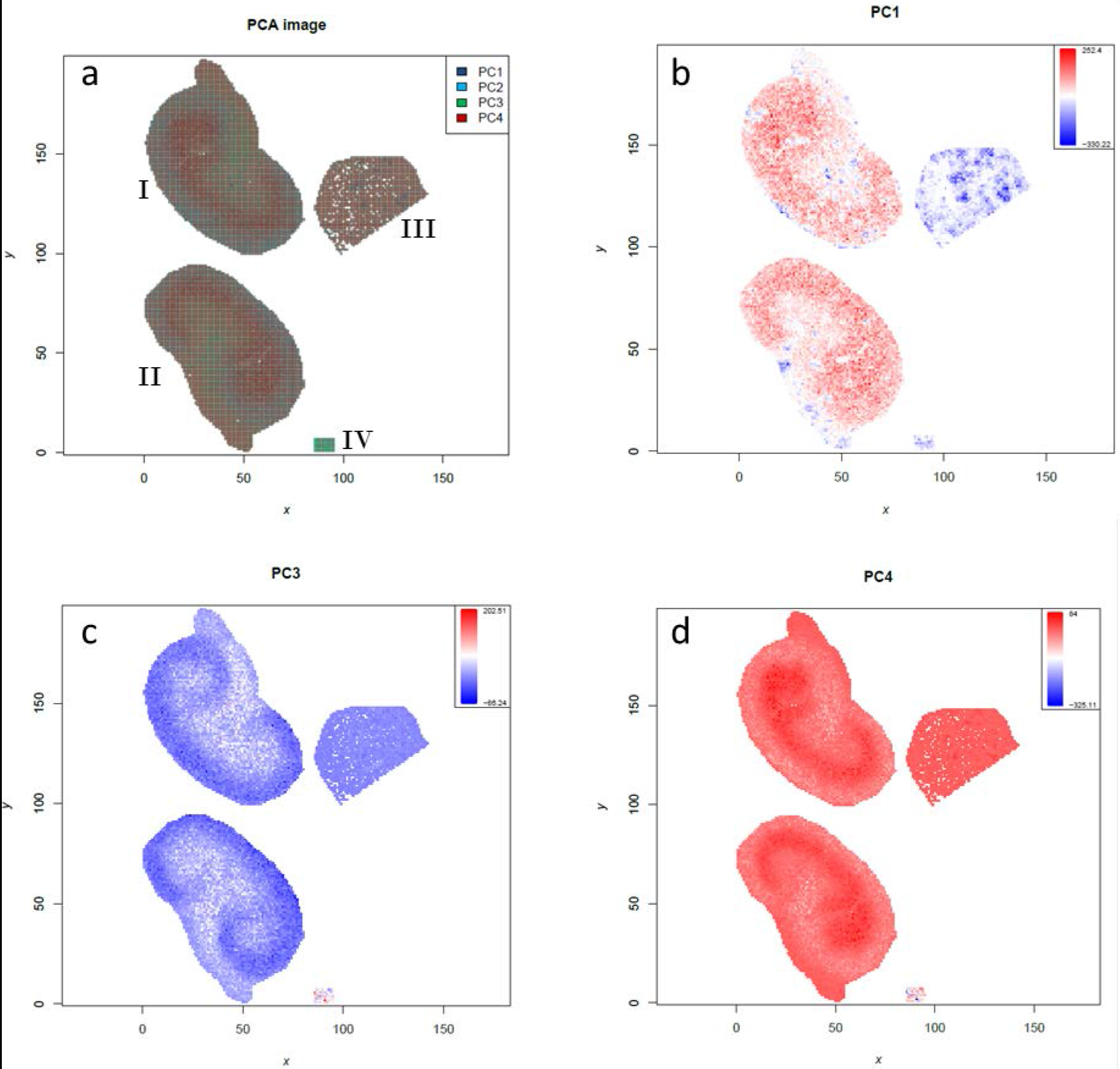
Results from the unsupervised statistical analysis of N-linked glycans in murine kidney tissues. Principal component analysis of treated, control and calibrant files. a) Overlay of all principal component scores. b-d) Principal components 1,3 and 4 that discriminate treated and control tissue or different kidney compartments. I = treated kidney1, II = treated kidney2, III = control kidney, IV = calibrants.

The complete analysis was performed in the European Galaxy instance with MSI tools based on Cardinal version 1.12.1 and MALDIquant 1.18 [21, 29]. The analysis history and workflow accompanies this publication as supporting information. Despite having used different algorithms for preprocessing and statistical analysis, we reached similar findings as compared to Gustafsson.

**Figure 3:**
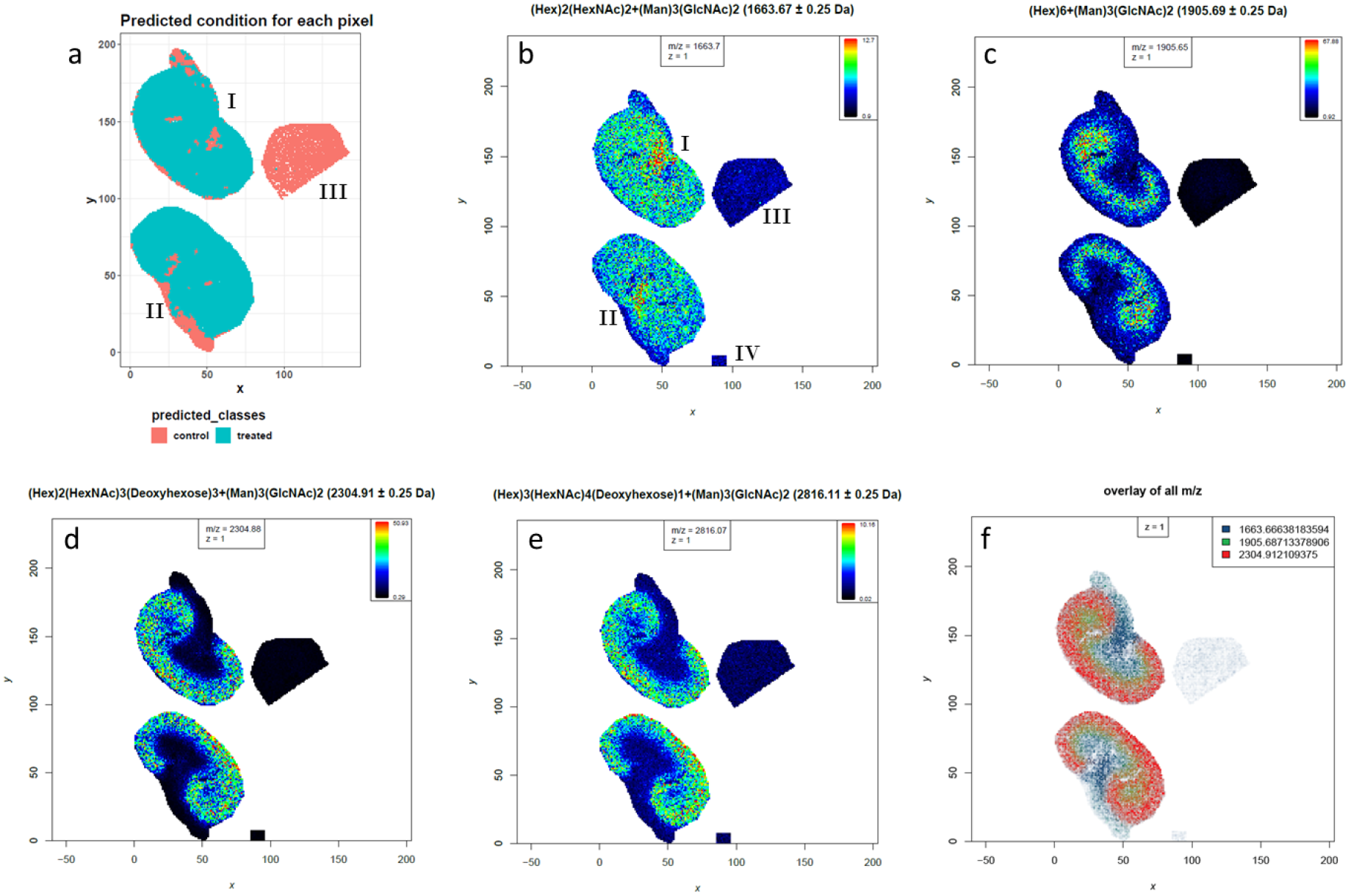
Results from the supervised statistical analysis of N-linked glycans in murine kidney tissues. The supervised spatial shrunken centroids method was used to determine m/z features that are more abundant in the treated than in the control tissue specimen. a) Spatial shrunken centroids class prediction for all spectra. b-e) Intensity distribution images for four identified N-linked glycans (Hex)_2_(HexNAc)_2_(Deoxyhexose)_1_+(Man)_3_(GlcNAc)_2_ (m/z 1663.6), (Hex)_6_+(Man)_3_(GlcNAc)_2_ (m/z 1905.7), (Hex)_2_(HexNAc)_3_(Deoxyhexose)_3_+(Man)_3_(GlcNAc)_2_ (m/z 2304.9) and (Hex)_3_(HexNAc)_4_(Deoxyhexose)_1_+(Man)_3_(GlcNAc)_2_(m/z 2816.0). f) Overlay of three N-linked glycans with different distribution in the kidney. The ion distribution images and the overlay image were generated with contrast enhancement by suppression on TIC normalized data. I = treated kidney1, II = treated kidney2, III = control kidney, IV = calibrants.

In Gustafsson’s own terms from a recent publication, our results show that their results are reproducible, because we, as another group, have followed as closely as possible their data analysis procedure and arrived at similar results [16].The reproducibility of the results shows the capacity of our pipeline. To enable what Gustafsson has described as “methods reproducibility” we provide the complete analysis history and the corresponding workflow. With this in hand, any other researcher can use the same tools and parameters in Galaxy to obtain the same result as we did.

**Table 1:** N-linked Glycans identified in the re-analysis. We could identify 16 N-linked glycans by matching the m/z features of the MSI data (column 1) to the identified m/z features of the LC-MS/MS experiment (column 5). We allowed a maximum tolerance of 300 ppm and multiple matches. Only single matches occurred with an average m/z error of 46 ppm (column 6).

| mz | centers | tstatistics | adj.p.values | M+Na+ | composition | ppm |
| --- | --- | --- | --- | --- | --- | --- |
| 1257.47424 | 38.24 | 51.97 | 0 | 1257.41 | (Hex)<sub>2</sub>+(Man)<sub>3</sub>(GlcNAc)<sub>2</sub> | 51 |
| 1743.68713 | 32.11 | 48.56 | 0 | 1743.57 | (Hex)<sub>5</sub>+(Man)<sub>3</sub>(GlcNAc)<sub>2</sub> | 67 |
| 1419.55334 | 40.68 | 48.2 | 0 | 1419.47 | (Hex)<sub>3</sub>+(Man)<sub>3</sub>(GlcNAc)<sub>2</sub> | 59 |
| 1905.68713 | 48.61 | 44.78 | 0 | 1905.63 | (Hex)<sub>6</sub>+(Man)<sub>3</sub>(GlcNAc)<sub>2</sub> | 30 |
| 2304.91211 | 43.53 | 42.36 | 0 | 2304.83 | (Hex)<sub>2</sub>(HexNAc)<sub>3</sub>(Deoxyhexose)<sub>3</sub>+(Man)<sub>3</sub>(GlcNAc)<sub>2</sub> | 36 |
| 1850.71216 | 25.3 | 42.01 | 0 | 1850.65 | (Hex)<sub>1</sub>(HexNAc)<sub>3</sub>(Deoxyhexose)<sub>1</sub>+(Man)<sub>3</sub>(GlcNAc)<sub>2</sub> | 34 |
| 1581.62573 | 18.07 | 40.64 | 0 | 1581.53 | (Hex)<sub>4</sub>+(Man)<sub>3</sub>(GlcNAc)<sub>2</sub> | 61 |
| 1809.72461 | 10.81 | 38.15 | 0 | 1809.63 | (Hex)<sub>2</sub>(HexNAc)<sub>2</sub>(Deoxyhexose)<sub>1</sub>+(Man)<sub>3</sub>(GlcNAc)<sub>2</sub> | 52 |
| 2158.88721 | 14.77 | 38.03 | 0 | 2158.77 | (Hex)<sub>2</sub>(HexNAc)<sub>3</sub>(Deoxyhexose)<sub>2</sub>+(Man)<sub>3</sub>(GlcNAc)<sub>2</sub> | 54 |
| 1663.66638 | 10.27 | 32.26 | 0 | 1663.57 | (Hex)<sub>2</sub>(HexNAc)<sub>2</sub>+(Man)<sub>3</sub>(GlcNAc)<sub>2</sub> | 58 |
| 1688.71509 | 8.68 | 28.29 | 0 | 1688.61 | (HexNAc)<sub>3</sub>(Deoxyhexose)<sub>1</sub>+(Man)<sub>3</sub>(GlcNAc)<sub>2</sub> | 62 |
| 1485.62378 | 8.67 | 26.89 | 0 | 1485.53 | (HexNAc)<sub>2</sub>(Deoxyhexose)<sub>1</sub>+(Man)<sub>3</sub>(GlcNAc)<sub>2</sub> | 63 |
| 2012.78394 | 7.3 | 26.72 | 0 | 2012.71 | (Hex)<sub>2</sub>(HexNAc)<sub>3</sub>(Deoxyhexose)<sub>1</sub>+(Man)<sub>3</sub>(GlcNAc)<sub>2</sub> | 37 |
| 2816.11206 | 6.92 | 26.35 | 0 | 2816.01 | (Hex)<sub>3</sub>(HexNAc)<sub>4</sub>(Deoxyhexose)<sub>1</sub>+(Man)<sub>3</sub>(GlcNAc)<sub>2</sub> | 36 |
| 2067.75903 | 5.69 | 14.52 | 0 | 2067.67 | (Hex)<sub>7</sub>+(Man)<sub>3</sub>(GlcNAc)<sub>2</sub> | 43 |
| 1542.61902 | 5.59 | 8.08 | 0 | 1542.55 | (HexNAc)<sub>3</sub>+(Man)<sub>3</sub>(GlcNAc)<sub>2</sub> | 45 |

Publishing histories and workflows from Galaxy requires only a few clicks and provides more information than requested by the minimum reporting guidelines MSI MIAPE (Minimum Information About a Proteomics Experiment) and MIAMSIE (Minimum Information About a Mass Spectrometry Imaging Experiment) [6, 16]. The Galaxy software itself but also the shared histories and workflows fulfil the FAIR principles that stand for findability, accessibility, interoperability, and reusability [27].

#### Summary

With the integration of the MSI data analysis toolset, we have incorporated an accessible and reproducible data analysis platform for MSI data in the Galaxy framework. Our MSI tools complement the multitude of already available Galaxy tools for proteomics and metabolomics that are maintained by Galaxy-M, Galaxy-P, Phenomal and Workflow4Metabolomics [34–38]. We are in close contact with those communities and would like to encourage developers of the MSI community to join forces and make their tools available in the Galaxy framework. We currently focused on reproducible and accessible data analysis, but we are planning to integrate interactive visualizations, more support for very large files and more tools for specific use cases into the Galaxy framework. Lastly, we would like to invite the MSI community to use the advantages of the Galaxy framework to advance MSI data analysis.

## Availability of supporting source code and requirements

Project name: Galaxy-P

Project homepage: https://github.com/galaxyproteomics/tools-galaxyp and https://github.com/BMCV/galaxy-image-analysis

Galaxy Toolshed: https://toolshed.g2.bx.psu.edu/

Operating system(s): Unix (Platform independent with Docker)

Training repository: https://galaxyproject.github.io/training-material/

Docker image: https://github.com/foellmelanie/docker-galaxy-msi

License: MIT

## Additional files

Additional file 1: Overview of R-functions in the MSI tools. For each Galaxy MSI tool the R-functions that do not belong to the basic R-package are listed.

## Availability of supporting data

Galaxy workflow to convert Bruker ROI.xml files: https://usegalaxy.eu/u/melanie-foell/w/msi-workflow-bruker-xml-conversion-to-tabular-file

Galaxy workflow to convert Bruker spotlists: https://usegalaxy.eu/u/melanie-foell/w/bruker-spotlist-conversion-to-tabular-file

Galaxy workflow co-registration: https://usegalaxy.eu/u/melanie-foell/w/co-registration-of-msi-image-and-real-image-with-landmarks

Galaxy workflow N-linked glycans re-analysis: https://usegalaxy.eu/u/melanie-foell/w/msi-workflow-complete-n-glycan-analysis

Galaxy history N-linked glycans re-analysis: https://usegalaxy.eu:/u/melanie-foell/h/re-analysis-of-pride-dataset-pxd009808---maldi-imaging-of-n-linked-glycans-in-murine-kidney-specimens

## List of abbreviations

DESI: Desorption Electrospray Ionization
MALDI: Matrix Assisted Laser Desorption/Ionization
FFPE: formalin-fixed paraffin-embedded
LC-MS/MS: liquid chromatography tandem mass spectrometry
MIAMSIE: Minimum Information About a Mass Spectrometry Imaging Experiment
MIAPE: minimum information about a proteomics experiment
MSI: Mass spectrometry imaging
PRIDE: proteomics identifications
ROI: Region of interest
SIMS: Secondary Ion Mass Spectrometry
TOF: Time of flight

## Authors’ contributions

M.C.F. developed the MSI Galaxy tool wrappers, the training material and the case study. L.M. acquired data for the training material, tested MSI tools and training material and provided useful feedback. T.W. developed the Galaxy tools and workflow for co-registration and contributed to build the Galaxy tool wrappers. M.N.S. tested MSI tools, co-registration tools and training material and provided useful feedback. N.V. built the Galaxy tool wrappers for the co-registration tools and tested them. M.W., P.B., K.R., B.A.G., O.S. contributed to the conceptualization, methodology, and funding acquisition. B.A.G. integrated the MSI file formats into Galaxy, contributed to build the training material and tool wrappers and integrated all tool wrappers into Galaxy. O.S. and M.F. wrote the manuscript. All authors critically read and approved the manuscript’s contents.

## Acknowledgement

We thank the European Galaxy Instance for bioinformatics support (https://usegalaxy.eu/) and the Galaxy community for critically reviewing tools and training material.

## References

1. Yang B, Patterson NH, Tsui T, Caprioli RM, Norris JL. Single-Cell Mass Spectrometry Reveals Changes in Lipid and Metabolite Expression in RAW 264.7 Cells upon Lipopolysaccharide Stimulation. J Am Soc Mass Spectrom. 2018;29:1012–20.

2. Bhandari DR, Wang Q, Friedt W, Spengler B, Gottwald S, Römpp A. High resolution mass spectrometry imaging of plant tissues: towards a plant metabolite atlas. Analyst. 2015;140:7696–709.

3. Bradshaw R, Bleay S, Clench MR, Francese S. Direct detection of blood in fingermarks by MALDI MS profiling and Imaging. Sci Justice. 2014;54:110–7.

4. Correa DN, Zacca JJ, Rocha WF de C, Borges R, de Souza W, Augusti R, et al. Anti-theft device staining on banknotes detected by mass spectrometry imaging. Forensic Sci Int. 2016;260:22–6.

5. Kramell AE, García-Altares M, Pötsch M, Kluge R, Rother A, Hause G, et al. Mapping Natural Dyes in Archeological Textiles by Imaging Mass Spectrometry. Sci Rep. 2019;9:2331.

6. McDonnell LA, Römpp A, Balluff B, Heeren RMA, Albar JP, Andrén PE, et al. Discussion point: Reporting guidelines for mass spectrometry imaging. Anal Bioanal Chem. 2015;407:2035–45.

7. Vaysse PM, Heeren RMA, Porta T, Balluff B. Mass spectrometry imaging for clinical research-latest developments, applications, and current limitations. Analyst. 2017;142:2690–712.

8. Karlsson O, Hanrieder J. Imaging mass spectrometry in drug development and toxicology. Arch Toxicol. 2017;91:2283–94.

9. Hoffmann WD, Jackson GP. Forensic Mass Spectrometry. Annu Rev Anal Chem. 2015;8:419–40.

10. Römpp A, Both JP, Brunelle A, Heeren RMA, Laprévote O, Prideaux B, et al. Mass spectrometry imaging of biological tissue: an approach for multicenter studies. Anal Bioanal Chem. 2015;407:2329–35.

11. Buck A, Heijs B, Beine B, Schepers J, Cassese A, Heeren RMA, et al. Round robin study of formalin-fixed paraffin-embedded tissues in mass spectrometry imaging. Anal Bioanal Chem. 2018;:1–12.

12. Porcari AM, Zhang J, Garza KY, Rodrigues-Peres RM, Lin JQ, Young JH, et al. Multicenter Study Using Desorption-Electrospray-Ionization-Mass-Spectrometry Imaging for Breast-Cancer Diagnosis. Anal Chem. 2018;90:11324–32.

13. Ly A, Longuespée R, Casadonte R, Wandernoth P, Schwamborn K, Bollwein C, et al. Site-to-Site Reproducibility and Spatial Resolution in MALDI-MSI of Peptides from Formalin-Fixed Paraffin-Embedded Samples. PROTEOMICS - Clin Appl. 2018;:1800029.

14. Gessel MM, Norris JL, Caprioli RM. MALDI imaging mass spectrometry: Spatial molecular analysis to enable a new age of discovery. J Proteomics. 2014;107:71–82.

15. Grüning B, Chilton J, Köster J, Dale R, Soranzo N, van den Beek M, et al. Practical Computational Reproducibility in the Life Sciences. Cell Syst. 2018;6:631–5.

16. Gustafsson OJR, Winderbaum LJ, Condina MR, Boughton BA, Hamilton BR, Undheim EAB, et al. Balancing sufficiency and impact in reporting standards for mass spectrometry imaging experiments. Gigascience. 2018;7:1–13.

17. Gruening B, Sallou O, Moreno P, da Veiga Leprevost F, Ménager H, Søndergaard D, et al. Recommendations for the packaging and containerizing of bioinformatics software. F1000Research. 2018;7:742.

18. Schramm T, Hester A, Klinkert I, Both JP, Heeren RM a, Brunelle A, et al. ImzML - A common data format for the flexible exchange and processing of mass spectrometry imaging data. J Proteomics. 2012;75:5106–10.

19. Gamboa-Becerra R, Ramírez-Chávez E, Molina-Torres J, Winkler R. MSI.R scripts reveal volatile and semi-volatile features in low-temperature plasma mass spectrometry imaging (LTP-MSI) of chilli (Capsicum annuum). Anal Bioanal Chem. 2015;:5673–84.

20. Gibb S, Strimmer K. Maldiquant: A versatile R package for the analysis of mass spectrometry data. Bioinformatics. 2012;28:2270–1.

21. Bemis KD, Harry A, Eberlin LS, Ferreira C, van de Ven SM, Mallick P, et al. Cardinal: an R package for statistical analysis of mass spectrometry-based imaging experiments. Bioinformatics. 2015;31:2418–20.

22. Veselkov K, Sleeman J, Claude E, Vissers JPC, Galea D, Mroz A, et al. BASIS: High-performance bioinformatics platform for processing of large-scale mass spectrometry imaging data in chemically augmented histology. Sci Rep. 2018;8:1–11.

23. Ràfols P, Torres S, Ramírez N, Del Castillo E, Yanes O, Brezmes J, et al. rMSI: an R package for MS imaging data handling and visualization. Bioinformatics. 2017;33:2427–8.

24. van der Walt S, Schönberger JL, Nunez-Iglesias J, Boulogne F, Warner JD, Yager N, et al. scikit-image: image processing in Python. PeerJ. 2014;2:e453.

25. Afgan E, Baker D, Batut B, Van Den Beek M, Bouvier D, Čech M, et al. The Galaxy platform for accessible, reproducible and collaborative biomedical analyses: 2018 update. Nucleic Acids Res. 2018;46:W537–-W544.

26. Taylor CF, Paton NW, Lilley KS, Binz P, Julian RK, Jones AR, et al. The minimum information about a proteomics experiment (MIAPE). Nat Biotechnol. 2007;25:887–93.

27. Wilkinson MD. Comment: The fair guiding principles for scientific data management and stewardship. Sci Data. 2016;:1–9.

28. Main Galaxy Instance. https://usegalaxy.org/. Accessed 2 Apr 2019.

29. European Galaxy Instance. https://usegalaxy.eu/. Accessed 9 Mar 2019.

30. Australian Galaxy Instance. https://usegalaxy.org.au/. Accessed 2 Apr 2019.

31. Grüning B, Dale R, Sjödin A, Chapman BA, Rowe J, Tomkins-Tinch CH, et al. Bioconda: sustainable and comprehensive software distribution for the life sciences. Nat Methods. 2018;15:475–6.

32. Boekel J, Chilton JM, Cooke IR, Horvatovich PL, Jagtap PD, Käll L, et al. Multi-omic data analysis using Galaxy. Nat Biotechnol. 2015;33:137–9.

33. Heydarian M. Prediction of Gene Activity in Early B Cell Development Based on an Integrative Multi-Omics Analysis. J Proteomics Bioinform. 2014;7.

34. Davidson RL, Weber RJM, Liu H, Sharma-Oates A, Viant MR. Galaxy-M: A Galaxy workflow for processing and analyzing direct infusion and liquid chromatography mass spectrometry-based metabolomics data. Gigascience. 2016;5.

35. Jagtap PD, Johnson JE, Onsongo G, Sadler FW, Murray K, Wang Y, et al. Flexible and Accessible Workflows for Improved Proteogenomic Analysis Using the Galaxy Framework. J Proteome Res. 2014;13:5898–908.

36. Peters K, Bradbury J, Bergmann S, Capuccini M, Cascante M, de Atauri P, et al. PhenoMeNal: processing and analysis of metabolomics data in the cloud. Gigascience. 2019;8:1–12.

37. Guitton Y, Tremblay-Franco M, Le Corguillé G, Martin JF, Pétéra M, Roger-Mele P, et al. Create, run, share, publish, and reference your LC-MS, FIA-MS, GC-MS, and NMR data analysis workflows with the Workflow4Metabolomics 3.0 Galaxy online infrastructure for metabolomics. Int J Biochem Cell Biol. 2017; January:1–13.

38. Galaxy-P Github repository. https://github.com/galaxyproteomics/tools-galaxyp. Accessed 2 Apr 2019.

39. da Veiga Leprevost F, Grüning BA, Alves Aflitos S, Röst HL, Uszkoreit J, Barsnes H, et al. BioContainers: an open-source and community-driven framework for software standardization. Bioinformatics. 2017;33:2580–2.

40. Mass Spectrometry Imaging Society: Software tools. https://ms-imaging.org/wp/imzml/software-tools/. Accessed 9 Mar 2019.

41. Grüning BA, Rasche E, Rebolledo-Jaramillo B, Eberhard C, Houwaart T, Chilton J, et al. Jupyter and Galaxy: Easing entry barriers into complex data analyses for biomedical researchers. PLoS Comput Biol. 2017;13:e1005425.

42. Wollmann T, Erfle H, Eils R, Rohr K, Gunkel M. Workflows for microscopy image analysis and cellular phenotyping. J Biotechnol. 2017;261 July:70–5.

43. GNU Image Manipulation Program (GIMP). https://www.gimp.org/. Accessed 2 Apr 2019.

44. Giacomoni F, Le Corguillé G, Monsoor M, Landi M, Pericard P, Pétéra M, et al. Workflow4Metabolomics: A collaborative research infrastructure for computational metabolomics. Bioinformatics. 2015;31:1493–5.

45. Bemis KD, Ferreira CR, Harry A, van de Ven SM, Mallick P, Stolowitz M, et al. Probabilistic Segmentation of Mass Spectrometry (MS) Images Helps Select Important Ions and Characterize Confidence in the Resulting Segments. Mol Cell Proteomics. 2016;15:1761–72.

46. Alexandrov T, Kobarg JH. Efficient spatial segmentation of large imaging mass spectrometry datasets with spatially aware clustering. Bioinformatics. 2011;27:230–8.

47. Bemis KD, Ferreira CR, Harry A, van de Ven SM, Mallick P, Stolowitz M, et al. Probabilistic Segmentation of Mass Spectrometry (MS) Images Helps Select Important Ions and Characterize Confidence in the Resulting Segments. Mol Cell Proteomics. 2016;15:1761–72.

48. UniProt Consortium. UniProt: a worldwide hub of protein knowledge. Nucleic Acids Res. 2019;47:D506–15.

49. Sud M, Fahy E, Cotter D, Brown A, Dennis EA, Glass CK, et al. LMSD: LIPID MAPS structure database. Nucleic Acids Res. 2007;35 Database:D527–32.

50. Blankenberg D, Von Kuster G, Bouvier E, Baker D, Afgan E, Stoler N, et al. Dissemination of scientific software with Galaxy ToolShed. Genome Biol. 2014;15:403.

51. Batut B, Hiltemann S, Bagnacani A, Baker D, Bhardwaj V, Blank C, et al. Community-Driven Data Analysis Training for Biology. Cell Syst. 2018;6:752–758.e1.

52. Galaxy Training Network. https://galaxyproject.github.io/training-material/. Accessed 9 Mar 2019.

53. Moritz L, Föll M, Schilling O. MALDI imaging of mouse kidney peptides - test dataset on Zenodo. 2018. doi:10.5281/zenodo.1560645.

54. Gustafsson OJR, Briggs MT, Condina MR, Winderbaum LJ, Pelzing M, McColl SR, et al. MALDI imaging mass spectrometry of N-linked glycans on formalin-fixed paraffin-embedded murine kidney. Anal Bioanal Chem. 2015;407:2127–39.

55. Gustafsson OJR, Briggs MT, Condina MR, Winderbaum LJ, Pelzing M, McColl SR, et al. Raw N-glycan mass spectrometry imaging data on formalin-fixed mouse kidney. Data Br. 2018;21:185–8.

56. Vizcaíno JA, Csordas A, Del-Toro N, Dianes JA, Griss J, Lavidas I, et al. 2016 update of the PRIDE database and its related tools. Nucleic Acids Res. 2016;44:D447–56.

57. Gibb S, Strimmer K. Mass Spectrometry Analysis Using MALDIquant. In: Statistical Analysis of Proteomics, Metabolomics, and Lipidomics Data Using Mass Spectrometry. Cham: Springer International Publishing; 2017. p. 101–24.

